# Chronic oral cannabidiol delays or prevents seizures in a mouse model of CLN2 disease

**DOI:** 10.1101/2025.11.17.688949

**Authors:** Joshua T. Dearborn, Keigo Takahashi, Nicholas R. Rensing, Michael Wong, Jonathan D. Cooper, Mark S. Sands

## Abstract

A growing body of literature describes the anti-inflammatory, neuroprotective, and anti-epileptic properties of the *cannabis sativa* constituent cannabidiol, suggesting that it might play a useful role in the treatment of neurodegenerative diseases. Late infantile neuronal ceroid lipofuscinosis (CLN2 disease) is a rare pediatric neurodegenerative disorder resulting from an inherited dysfunction of the lysosome. CLN2 disease, and its representative animal models, display neuroimmune response, neuroinflammation, neurodegeneration, and epileptic seizures, and these symptoms are all touted as potential targets of cannabidiol therapeutic benefit. Here, we treated a valid model of CLN2 disease with long-term daily cannabidiol (300mg/kg) from 1 month of age until disease end stage and evaluated epileptic seizures, lifespan, and markers of neuroimmune response. Chronic cannabidiol treatment significantly delayed or fully eliminated seizures in CLN2 mice compared to those treated with vehicle only, and the treatment led to a nearly significant extension of lifespan. These effects occurred in the absence of any therapeutic benefit to physiological markers of disease such as GFAP, CD68, and cytokine/chemokine reactivity. Taken together, we show that chronic treatment with cannabidiol confers significant anti-seizure benefit to the mouse model of CLN2 disease, and that it does not appear to do so by altering the inflammatory and neuroimmune markers traditionally used to track CLN2 disease progression.

## Introduction

Mutations in the *TPP1/CLN2* gene encoding the lysosomal enzyme tripeptidyl peptidase 1 (TPP1) cause CLN2 disease or late infantile Neuronal Ceroid Lipofuscinosis, previously referred to as classical late infantile Batten disease(1). Children with CLN2 disease experience developmental delay, epilepsy, and early death(2,3). There is currently one FDA-approved therapy for CLN2 disease, which involves regularly supplying recombinant TPP1 enzyme into the lateral ventricle via a chronic port. This treatment (Brineura enzyme replacement therapy) slows the progression of disease and improves some clinical symptoms(4). It must be supplied for the life of the patient and is administered in two-week increments. Despite some benefit, enzyme replacement therapy does not completely prevent or reverse progression of the disease, so most patients continue to have impairing neurological symptoms. The need for developing novel treatments for CLN2 disease is clear.

In recent years, purified cannabidiol (CBD; drug name Epidiolex) from the *cannabis sativa* plant has been approved for the treatment of seizures in children with Dravet Syndrome, Lennox-Gastaut Syndrome, and Tuberous Sclerosis Complex. Addition of CBD to existing anti-seizure drug regimens leads to an additional reduction in seizure frequency in children with these conditions(5–7). Anecdotally, some families of NCL patients report that various cannabis products help control their children’s seizures. However, there have been no controlled studies in children with CLN2 disease or animal models of the disease. There are many reports of other effects of CBD in animal models of disease. Germane to CLN2, it has been shown that CBD may have anti-inflammatory, neuroimmunomodulatory, and neuroprotective effects(8–12). Our own work evaluated the effects of chronic CBD therapy in the mouse model (*Cln1*^*-/-*^) of a related disorder, CLN1 disease/infantile NCL, previously referred to as infantile Batten disease. We showed that chronic oral administration of CBD reduced the neuroimmune response in areas of the thalamus and cortex, well-established as being severely affected regions of the CNS(13,14).

CBD appears to have limited neuroprotective effects as measured by neuron count and cortical thickness, however these measures were taken at CLN1 disease end-stage. There also appeared to be no effect of CBD on seizure frequency in the *Cln1*^*-/-*^ mouse. Importantly, based on existing literature, the dose of CBD that was administered to the *Cln1*^*-/-*^ mouse was relatively low compared to those used in other disorders (11,15–17). It is our belief that a higher dose of CBD may yield a more positive therapeutic benefit.

A mouse model of CLN2 disease harboring the *Cln2*^*R207X*^ mutation displays some important similarities to the *Cln1*^*-/-*^ mouse we previously characterized. It shows many of the same phenotypes including neuroimmune response, neurodegeneration, and spontaneously arising seizures(14,18,19). Of particular interest is that the seizure phenotype observed in CLN2 mice is more profound than that seen in *Cln1*^*-/-*^ mice; the onset of seizures is earlier and the CLN2 mice typically die within minutes of a seizure(20). In addition, though there are some important differences in onset and severity of the pathology, *Cln2*^*R207X*^ mice show microglial and astrocyte activation and neuron loss like that seen in the *Cln1*^*-/-*^ mouse in the same central regions. Just like in the *Cln1*^*-/-*^ mouse, the pathogenesis of seizures in the *Cln2*^*R207X*^ remains unknown.

Given the limited effects of the 100mg/kg dose on the neuroimmune response and no relief of seizures in the *Cln1*^*-/-*^ mouse, we chose a higher dose (300mg/kg) for the CLN2 mouse. This dose falls within the range of those used in other preclinical seizures model(11,12,15,16,21). We reasoned that a relatively high dose of CBD would be necessary for this more rapidly progressing disease model with more severe seizures. We evaluated the effects of chronic CBD therapy on glial activation, disease-associated pathology, seizure frequency, and lifespan in the *Cln2*^*R207X*^ mouse model of CLN2 disease. We found that, even in the absence of improvements to any known CLN2-related pathophysiology, chronic treatment with 300mg/kg CBD significantly delays or prevents seizures and nearly significantly extends the lifespan of the mouse.

## Methods

### Animals

The *Cln2*^*R207X*^ mouse was first generated as described by Geraets et al.(22). For this study, mice were generated from in-house crosses of heterozygous *Cln2*^*R207X*^ mice as detailed previously(23). All mice were housed in an animal facility at Washington University School of Medicine under a 12h light/12h dark cycle and had access to food and water *ad libitum*. Seizure activity was monitored by continuous video/EEG recordings in a total of 31 mice, both male and female, all *Cln2*^*R207X*^. It was our aim to evaluate an even number of mice in each treatment group, but limitations on animal colony survival and apparatus availability led to a slightly uneven distribution. For seizure monitoring, 16 CBD-treated and 15 vehicle treated mice completed evaluation; these mice were all used for lifespan data as well. A separate cohort of mice was used to harvest tissues for various pathological and biochemical assays. Of this second cohort, immunostaining was performed on 20 *Cln2*^*R207X*^ brain hemispheres (10 CBD-treated and 10 vehicle-treated) as well as 5 WT mice, while cytokines were measured in 26 brain hemispheres (8 *Cln2*^*R207X*^ mice treated with CBD, 8 *Cln2*^*R207X*^ mice treated with vehicle, 5 WT mice treated with CBD, and 5 WT mice treated with vehicle. The concentration of CBD was measured by mass spectrometry on brain homogenates from 8 *Cln2*^*R207X*^ animals, 4 of which were treated with CBD while the other 4 were treated with vehicle only.

### CBD administration

Cannabidiol was prepared in flavored gelatin cubes that were provided to the mice for voluntary oral consumption. Our own previous work(13) details preparation of the gelatin cubes, though here they were made to contain either CBD (300mg/kg) in ethanol, or ethanol only. It was confirmed that across 14 consecutive days, mice reliably consume 60-75% of the gelatin cube (by weight; data not shown). CBD was dosed accordingly such that this proportion of a cube would contain the intended dose. Gelatin was provided to mice just prior to the onset of the dark cycle each day, with any remaining remnants replaced by a fresh cube the following day at the same time. This occurred every day from the age of 1 month until death (in the case of the EEG-monitored mice), or until the age of 3 months (for all other measures).

To confirm delivery of pure CBD to the brain via voluntary oral administration, brains of mice were analyzed via mass spectrometry following 2 months of daily CBD access, as detailed below.

### Tissue collection

After 2 months of treatment with either CBD or vehicle, mice were euthanized via *i*.*p*. injection of Fatal-Plus (Vortech Pharmaceuticals, Dearborn, MI, USA) and then perfused with PBS. The brain was removed, and one hemisphere was immediately flash-frozen in liquid nitrogen (for mass spectrometry and/or cytokine analysis) while the other was submerged in a solution of 4% paraformaldehyde in PBS for 48h to prepare for histological evaluation. After this 48h, preserved brain hemispheres were transferred to 30% sucrose in 40mM TBS (pH 7.6) until processing.

### Immunostaining

Sections of fixed brain tissue were stained on slides using a modified immunofluorescence protocol(18) for astrocytes (rabbit anti-GFAP, 1:1000, Agilent Z0334) and microglia (rat anti-mouse CD68, 1:400, Bio-Rad MCA1957) essentially as described previously (3). Coronal sections of 40µm were cut via Microm HM430 freezing microtome (Microm International GmbH, Wallendorf, Germany) and mounted on Superfrost plus slides (Fisher Scientific). After air-drying for 30min, slides were blocked in 15% serum solution (Normal goat serum, S-1000 Vector Laboratories) in 2% TBS-T (1 x Tris Buffered Saline, pH 7.6 with 2% Triton-X100, Fisher Scientific) for 1h. Slides were incubated in primary antibody in a 10% serum solution in 2% TBS-T for 2h, after which they were washed three times with 1xTBS. Slides were then incubated in fluorescent Alexa-Fluor-labelled IgG secondary antibodies (Alexa-Fluor goat anti-rabbit 488 Invitrogen A-11008, goat anti-rat 546 Invitrogen A-11081, both at 1:400 dilution) in 10% serum in 2% TBS-T for 2h. Another three 1xTBS washes were applied to the slides before they were incubated in a 1x solution of TrueBlack lipofuscin autofluorescence quencher (Biotium, Fremont, CA) in 70% ethanol for 2 min each. Slides were rinsed again with 1xTBS and coverslipped in fluoromount-G mounting medium with DAPI (Southern Biotech, Birmingham, AL).

### Thresholding image analysis

A semi-automated thresholding image analysis method(18) was performed to quantify glial activation in specific areas of interest. The ventral posteromedial/posterolateral (VPM/VPL) nuclei of the thalamus and the somatosensory barrel field (S1BF) were demarcated(24) after slides were scanned at 10x magnification. A one-in-six series of sections were captured per animal and lamp intensity, camera setup, and calibration were kept constant throughout image capture. Collected images were then analyzed using Image-Pro Premier (Media Cybernetics) to set a threshold that selected foreground immunoreactivity above background. This threshold was applied as a constant to all images for each region of interest and specific to the reagent used to detect immunoreactivity for each antigen.

### Mass spectrometry

Frozen brain tissue was analyzed in triplicate via liquid chromatography-tandem mass spectrometry (LC-MS/MS) as previously described (3) with minor modifications. Four µL injections of each sample were separated on a C18 column eluted at 0.25 mL/min with H_2_O with 0.1% formic acid (A) and acetonitrile with 0.1% formic acid (B) using the following gradient: 50% B (0-2 min), 50% B to 100% B (2-10 min), 100% B (15-16 min), and reequilibrated at 50% B (16-20 min). Sample ionization utilized a heated ESI source with the following settings: Spray voltage +3800 v, sheath gas 50 (arb), auxiliary gas 5 (arb), sweep gas 2 (arb), ion transfer tube 325 °C, and vaporizer temperature 200 °C. CBD and d_3_-CBD (IS) were detected using SRM (selected reaction monitoring). For CBD, the precursor ion was 315.2 m/z, and 259.21 m/z (collision energy 28.38 V) and 193.16 m/z (collision energy 32.34 V) were the product ions. For d_3_-CBD, the precursor ion was 318.3 m/z, and 262.21 m/z (collision energy 29.35 V) and 196.20 (collision energy 34.32 V) were the product ions. The SRM properties were as follows: Cycle time 0.8 s, Q1 resolution 0.7 FWHM, Q3 resolution 1.2 FWHM, CID gas (N_2_) 1.5 mTorr, and source fragmentation 0.

### Cytokine assays

Measures of chemokines and cytokines in brain homogenates were generated using a 12-biomarker Multi-Analyte Profile (MYRIAD RBM) using standard Luminex technology. The method has been previously described(13,25) and includes the following analytes: IL-10, IL-1β, IP-10, IL-4, IL-5, IL-6, IFN-γ, IL-12p70, GRO-α, TNF-α, MCP-1, and MIP-1β.

### Electroencephalography (EEG) electrode surgery

*Cln2*^*R207X*^ mice were implanted with EEG electrodes at postnatal day (PND) 53 essentially as described previously (13). Briefly, mice were placed under general anesthesia and kept warm while a midline vertical incision exposed the skull. The skull was cleaned and dried, then burr holes were made (anterio+0.5mm, lateral±0.5mm; bregma) using a micro drill before screws were secured to the skull and reference electrodes placed. Two bilateral “active” recording electrodes were placed over the parietal cortex (posterior -2.5mm, lateral±1.5mm; bregma), then a ground screw was placed over the cerebellum (posterior -6.2mm, lateral±0.5mm; bregma) via the same method as above. Dental cement (SNAP, Parkell) was applied to the exposed skull, screw, and wires to secure the pin header for subsequent recording. The skin was sutured, and tissue glue (Vetbond, 3M) closed the remainder of the incision, then mice received Buprenorphine (0.1mg/kg) while recovering in a warming chamber. After recovery, mice were placed in individual recording cages(13,26).

### Video-EEG monitoring

Mice were allowed to recover for at least 72h before simultaneous video/EEG monitoring began. A custom flexible cable was attached to the exposed pin header, and bilateral cortical EEG signals were collected using a referential montage via Stellate or LabChart (ADInstruments) acquisition software and amplifiers. As previously described (13,26), EEG signals were amplified, filtered, digitized, time-locked to video, and collected continuously until death or euthanasia. Electrographic seizures were identified by their characteristic pattern of discrete periods of rhythmic spike discharges that evolved in frequency and amplitude lasting at least 10 seconds, typically ending with repetitive burst discharges and voltage suppression. These instances were matched against corresponding video to confirm the spontaneous seizure behavioral phenotype (tail stiffening, abnormal rearing posture, “popcorn-like” and often violent propulsion about the chamber followed by stiffened immobility). A total of 31 *Cln2*^*R207X*^ mice (16 males, 15 females) were continuously monitored for EEG activity starting at PND 60, or approximately 2 months of age. Outcome variables include seizure occurrence, seizure duration, time between seizure and death, and lifespan.

### Statistical analysis

Data was analyzed using IBM SPSS Statistics (v28) software as well as GraphPad Prism 9. Where assumptions of normality were met, groups were compared using a Student’s *t-*test, and when data was distributed non-normally the nonparametric Mann-Whitney *U*-test was employed. A log rank Mantel-Cox analysis was used to evaluate survival. A *p* value < 0.05 was considered statistically significant. Data are expressed as mean ± standard error of the mean (SEM) except where indicated otherwise.

## Results

### CBD administration

Following two months of oral CBD administration, brain tissue from the four treated CLN2 mice contained an average of 28.53 ± 7.13ng of CBD per gram of brain tissue at time of euthanasia. The amount of CBD was below the level of quantification (LOQ) in the four vehicle-treated mice. Similar to a previous study in the CLN1 mouse (13), mass spectrometry confirmed that voluntary oral consumption of CBD via gelatin cube delivers measurable levels of drug to the brain.

### Neuroimmune and neuroinflammatory responses

Previously, immunohistology performed on untreated *Cln2*^*R207X*^ mouse brain tissue revealed significantly increased GFAP and CD68 signal throughout the cortex and in the thalamus(22,23). Similarly, in the current study, GFAP and CD68 staining in vehicle-treated *Cln2*^*R207X*^ mouse S1BF and VPM/VPL were significantly elevated compared to that seen in untreated WT mice. Chronic treatment with cannabidiol did not reduce the levels of either glial marker of neuroimmune response (Figure 1). We analyzed cytokines and chemokines in CBD- and vehicle-treated mouse brains to survey for inflammatory effects (Figure 2). Broadly speaking, these markers can be grouped into three categories: anti-inflammatory cytokines (IL-10, IL-4, and IL-5), monocyte activators (IP-10, MCP-1, and MIP-1β), and pro-inflammatory cytokines (IL-1β, IL-6, IFN-γ, IL12p70, GRO-α, and TNF-α)(25). In *Cln2*^*R207X*^ brain tissue exposed to CBD for 2 months, analysis revealed significantly increased levels of IL-1β compared to *Cln2*^*R207X*^ mice treated with vehicle only. There were no other differences between CBD-treated and vehicle-treated *Cln2*^*R207X*^ tissue on the cytokine/chemokine panel, and CBD treatment did not lead to any effect on levels measured in WT mice. Like in our previous findings(23), vehicle-treated *Cln2*^*R207X*^ mice did not differ from vehicle-treated WT mice on levels of the pro-inflammatory cytokines IL-1β, IL-6, and GROα, whereas here the CBD-treated *Cln2*^*R207X*^ mice showed increased levels of each compared to the same group (*p* = .004, *p* = .021, and *p* = .042 respectively).

**Figure 1.**
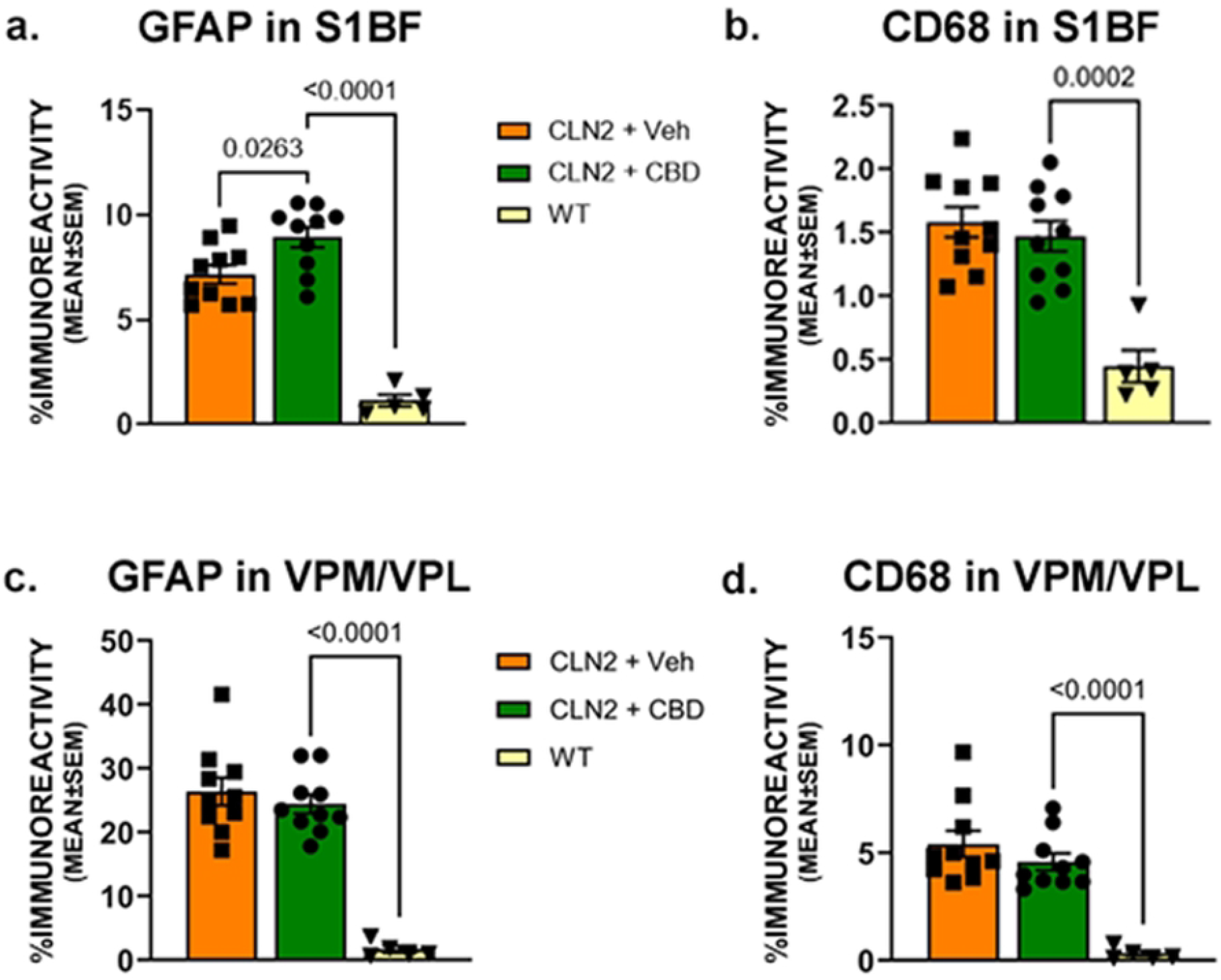

**Figure 2.**
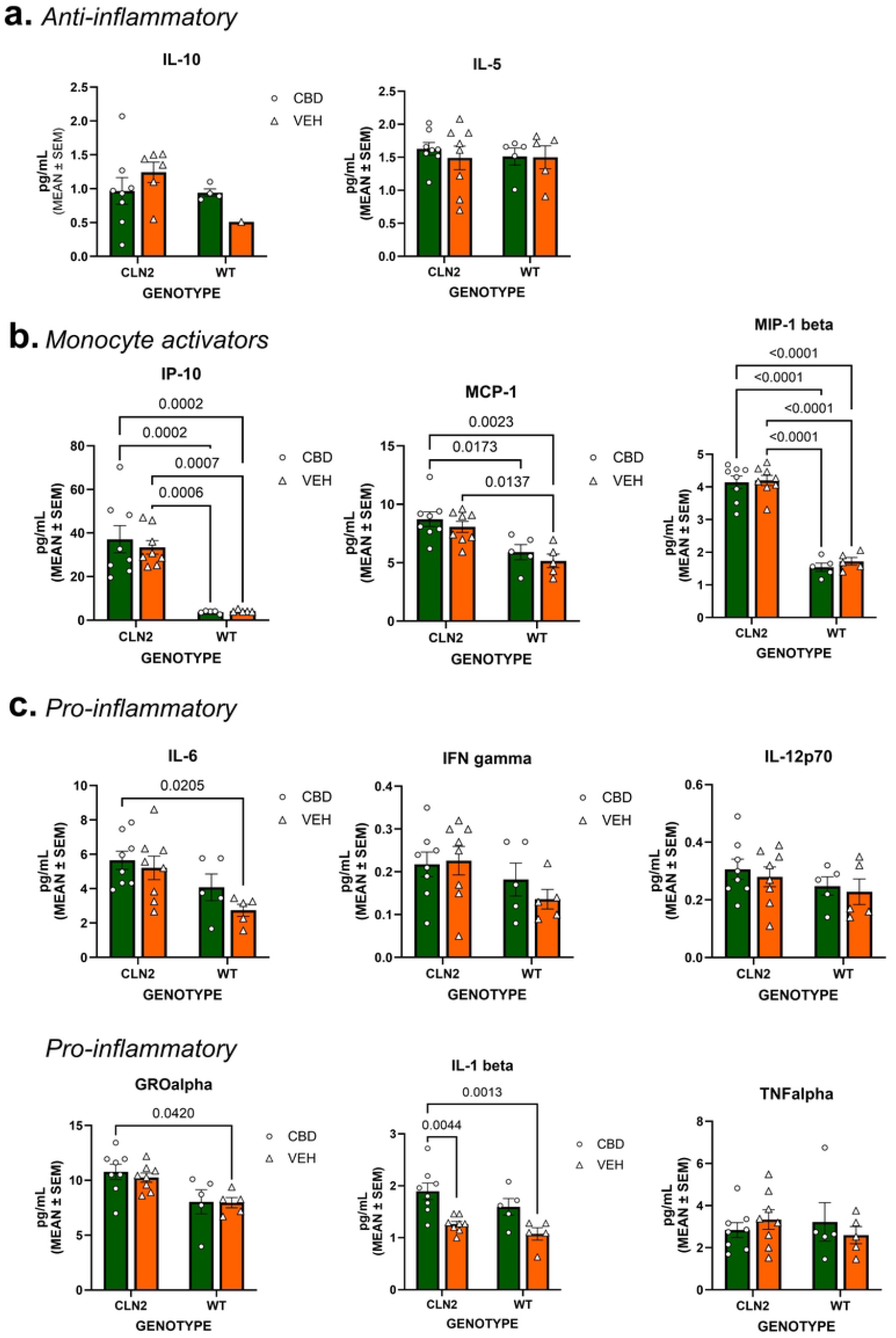

### Seizure activity

Chronic treatment with CBD significantly delayed or prevented the occurrence of a first seizure in *Cln2*^*R207X*^ mice [log-rank (Mantel-Cox) test, *p* = 0.033; Figure 3a]. Median age of first seizure in CBD-treated mice was 129 days, while median age of first seizure in vehicle-treated mice occurred at 113 days. A graphic depiction of seizure occurrence across lifespan for each mouse can be seen in Figure 3b. Though treatment with cannabidiol did not reduce the number of seizures per mouse (Mann-Whitney *U* = 77.50, *p* = 0.327) there was a trend towards a reduced mean duration as measured by EEG activity (*t* = 1.870, *df* = 79, *p* = 0.065; Figure 3c). When considering all mice, the mean duration of a single seizure per mouse was reduced by treatment with CBD (Mann-Whitney *U* = 68.50, *p* = 0.038; Figure 3d); when considering only mice that experienced at least one seizure, excluding any mice that did not, there was a trend toward reduced mean seizure duration in CBD-treated mice (*t* = 1.890, *df* = 19, *p* = 0.074; Figure 3e).

**Figure 3.**
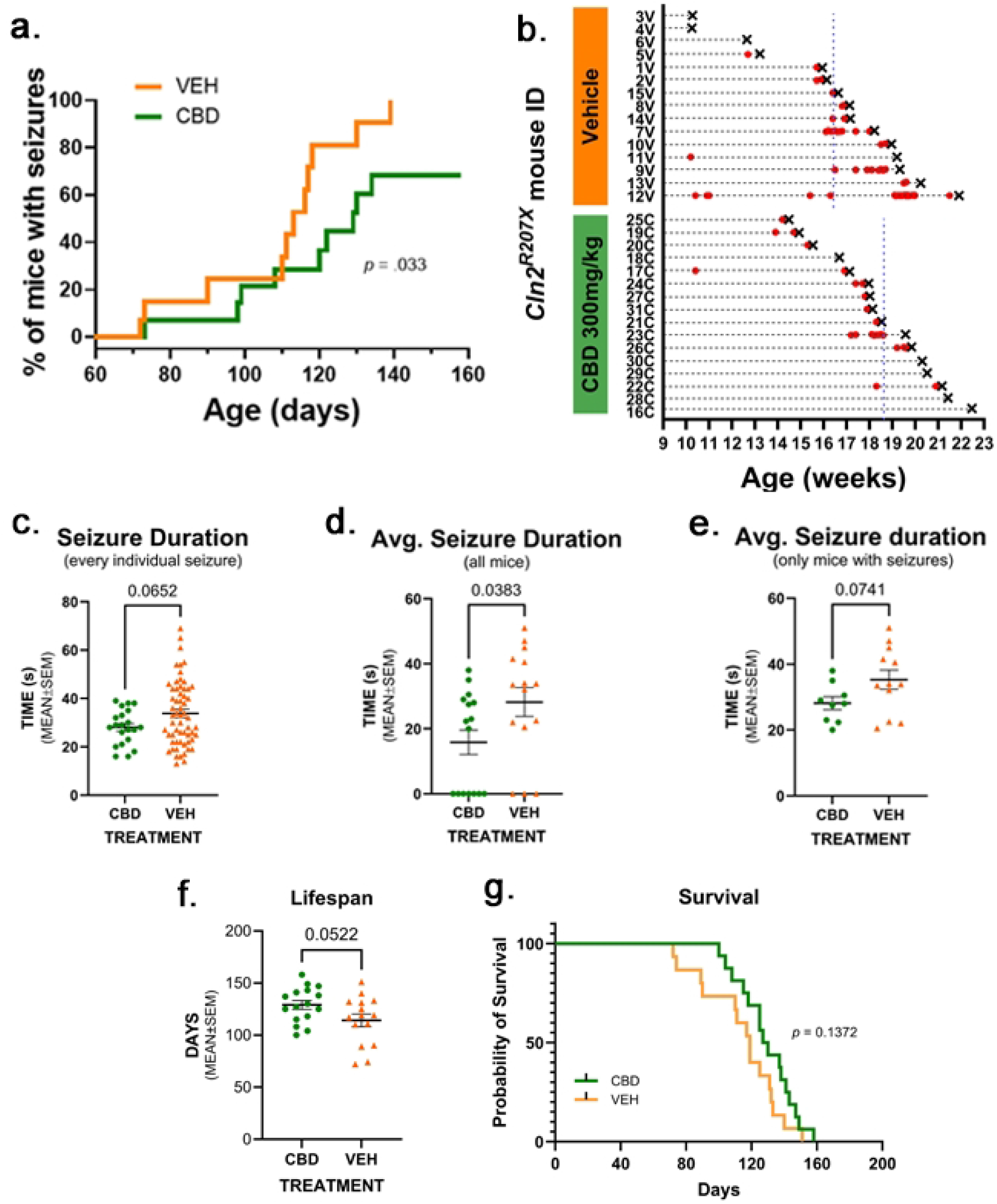

### Lifespan

*Cln2*^*R207X*^ mice treated with CBD lived longer than *Cln2*^*R207X*^ mice treated with vehicle alone, but this lifespan extension did not quite reach statistical significance (*t* = 2.025, *df* = 29, *p* = 0.052; Figure 3f) when measured by numbers of days survived; a log-rank (Mantel-Cox) test revealed no such trend (*p* = 0.137; Figure 3g) owing largely to a single overlapping data point in the longest-lived mice.

## Discussion

The purpose of this research was to evaluate whether chronic cannabidiol administration improved well defined pathology and seizure phenotypes in the *Cln2*^*R207X*^ mouse model of CLN2 disease. This rodent model recapitulates many facets of the human condition including CNS neuroinflammation and neuroimmune response, severe and fatal seizures, and a dramatically shortened lifespan. We chose an oral route of administration in order to more closely mimic how CBD is administered to patients, with the additional benefit that this delivery method is essentially ‘hands-off’. Animal stress is a critical variable since *Cln2*^*R207X*^ mice are very reactive to any perturbation. Anecdotally, entire cages of *Cln2*^*R207X*^ mice can die simply by being transferred between buildings via standard and accepted methods. We have also noticed that these mice are over-reactive to handling and often experience a fatal seizure as a result. It is known that seizures frequently lead to death in *Cln2*^*R207X*^ mice(20), and the gelatin cube method we employed here aided us in avoiding, or minimizing the risk of, this fate. We acknowledge that relying on the mice to voluntarily consume a flavored gelatin cube containing drug does not guarantee strict control over total dosage and timing. However, we found that most mice reliably consume 60-75% of the cubes over a 24h span(13). Most importantly, we confirmed delivery of CBD via this method to the brain of *Cln2*^*R207X*^ mice via mass spectrometry.

It is important to note here that CBD was administered in the absence of any other treatment that addressed the underlying genetic cause (ERT or gene therapy) or any other supportive care (i.e. anti-seizure or anti-inflammatory drugs). Here we show that while long-term treatment with CBD alone did not show a beneficial effect on neuroinflammation or neuroimmune response, it does delay or prevent seizures while nearly having a significant impact upon the lifespan of *Cln2*^*R207X*^ mice. The delayed onset or prevention of seizures, coupled with a nearly increased lifespan, is a remarkable finding given that the treatment appears not to have affected the well-described CNS pathology associated with CLN2 disease. Given what little is known about CBD’s precise mechanisms of actions, there is no reason to believe that the drug would address the underlying cause of CLN2 disease, TPP1 deficiency(1). It is not entirely unexpected, then, that CNS CLN2 pathology would be unaffected by a treatment that does not correct the underlying cause. Yet there is much evidence that CBD can reduce neuroinflammation in some circumstances, and that it can reduce seizures via thus far unknown mechanisms(5–12). Outside of the CNS, recent evidence reveals life-limiting disease pathology in the enteric nervous system (ENS), including the bowel (25,27–29). Given the oral route of administration we employed in the current study, it would follow that CBD might greatly impact the ENS leading to improved quality, and duration, of life. Regardless the mechanism, we have provided more evidence that chronic oral delivery of CBD alone can have a positive effect on seizures in a murine model of a particularly devastating neurodegenerative lysosomal storage disorder. In this way, it may be ripe for use as an adjunctive therapy, a treatment added to enzyme replacement therapy or the much-studied AAV-mediated gene therapy.

CBD has been widely touted as having antiseizure and anti-inflammatory properties. The *Cln2*^*R207X*^ mouse develops seizures naturally and as a consequence of known disease progression(20). Accordingly, this model system provides an authentic test of the symptom as compared to publications touting antiseizure effects in *inducible* models(8–12). In this mouse model of severe disease, chronic treatment with the drug significantly delayed or prevented development of seizures. Comparison via log-rank statistical analysis revealed that daily 300mg/kg cannabidiol delayed seizures in *Cln2*^*R207X*^ mice by 16 days compared to vehicle-treated counterparts. Importantly, these 16 days account for nearly 15% of the average lifespan of vehicle-treated *Cln2*^*R207X*^ mice in this study. It is notable that 5 of 16 CBD-treated mice never developed seizures, and the 3 vehicle-treated mice that were seizure-free died quite early compared to expected lifespan. Relatedly, CBD treatment nearly led to the CBD-treated mice living significantly longer than the vehicle-treated mice. The increase in lifespan of 15 days amounts to a 13% increase over the vehicle-treated *Cln2*^*R207X*^ mice. Again, this is remarkable based on the fact that neither the primary enzyme deficiency nor the profound changes in chemokines and cytokines or the neuroimmune response were treated.

We probed known CNS disease pathology for treatment effects, and to seek possible mechanisms for any effect CBD might have on the *Cln2*^*R207X*^ mouse. We surveyed immunohistology evidence of neuroimmune response, including the astrocytosis marker GFAP and the activated macrophage marker CD68, which are invariably elevated in the *Cln2*^*R207X*^ mouse brain(20,22). Specifically, the VPM/VPL of the thalamus and the S1BF of the cortex show significantly increased GFAP and CD68 immunoreactivity, respectively, in the CLN2 mouse model at 3 months of age(20). Here we recapitulated that finding and also report that chronic CBD treatment did not affect these measures at 3 months of age, when these mice are very severely affected. Our findings seem to indicate that: 1) chronic 300mg/kg of CBD does not improve measures of neuroimmune response known to be elevated in CLN2 disease, and 2) the positive effects on seizure and lifespan conferred by CBD are unlikely to be due to any effect on the neuroimmune response as measured here.

To evaluate disease-related neuroinflammation, we measured levels of 12 markers of cytokines and chemokines in brain tissue (Fig. 3). Generally speaking, these can be grouped into three categories: anti-inflammatory, pro-inflammatory, and monocyte activator. Brains from *Cln2*^*R207X*^ mice treated with vehicle exhibited significantly increased levels of the monocyte activators IP-10, MCP-1, and MIP-1β compared to brains from WT mice treated with vehicle only. This is unsurprising given the known effects of the disease on some markers of the immune system(20). Treatment with 300mg/kg CBD led to neither a reduction in any pro-inflammatory marker, nor an increase in any anti-inflammatory marker compared to the vehicle-treated mice. In fact, in *Cln2*^*R207X*^ mice, CBD treatment led to a statistically significant increase in the pro-inflammatory markers IL-1β, IL-6, and GROα (known also as CXCL1) compared to the vehicle group. It is worth nothing that while IL-6 is purported to be a pro-inflammatory marker in the central nervous system, it is suggested to play an anti-inflammatory role in the periphery, and specifically in muscles(30). As noted above, parts of the digestive system are significantly affected by CLN2 disease pathology. An in-depth survey of the enteric system would reveal the effects of cannabidiol on significant inflammation in a crucial system in the periphery. In the current study, while CBD exerted anti-seizure effects in the *Cln2*^*R207X*^ mice, it may not have done so via the neuroinflammation markers we surveyed, or it may have done so in a way that was antithetical to expectations.

Interestingly, while our previous work showed that chronic CBD reduced neuroimmune response in the CLN1 mouse, we did not find the same here. This difference might be viewed as somewhat of a conundrum in that the dose that was given to the CLN1 mouse was considerably lower than the one we used here (100mg/kg vs. 300mg/kg). However, it is important to note that although CLN1 disease and CLN2 disease are both within the family of NCL diseases, their differences are significant. The enzymes that are deficient in each disease, PPT1 in CLN1 and TPP1 in CLN2, have very different substrate specificities. PPT1 removes fatty acyl chains from palmitoylated proteins while TPP1 removes a unit of three amino acids from the amino terminus of proteins(31). As a result, accumulating substrates and altered organelle functions are different between CLN1 and CLN2 disease, likely resulting in different cellular consequences. Perhaps not surprisingly, the progression of disease pathology also appears to follow a different pattern in these mouse models. Pathology is first detectable in the spinal cord of CLN1 mice, then progresses to the thalamus and later the cortical areas. In contrast, the pathology in the CLN2 mouse is first detectable in the cortex, then later in the thalamus and spinal cord. It is also worth noting that the nature and timing of neuroimmune response and neuron loss differs between the two models (14,20,32–34) and that disease progression, based on lifespan and outwardly obvious symptoms, seems to be more rapid and severe in CLN2 mice. Given these patterns, it may not be surprising that the two mouse models of disease respond differently to CBD administration.

Use of CBD as therapy for a litany of conditions continues to spread, and a growing body of literature supports its effectiveness even in the absence of distinct mechanisms. Similarly, we were unable to determine the mechanism by which CBD exerted its positive effects on the CLN2 mouse. However, chronic oral administration of CBD conferred anti-seizure and life-extending benefits to the *Cln2*^*R207X*^ mice, perhaps due to treating disease outside of the CNS. Clearly more sensitive or comprehensive methods must be employed to determine the mechanism of CBD action in this model system. This experiment was not designed as a dose-response study. Rather, we chose a dose of CBD that is comparable to what is used clinically. Based on the half-life of CBD in humans and mice(35), the dose we provided our CLN2 mice (300mg/kg) is roughly equivalent to 10mg/kg in humans, falling well within the range used clinically. It is certainly possible that a higher dose of CBD or earlier initiation of treatment would have resulted in greater therapeutic efficacy. Alternatively, CBD might synergize with other anti-seizure drugs or with a therapy that specifically targets the underlying genetic cause.

## Conclusions

Taken together, we provide evidence that chronic cannabidiol can protect against seizures and nearly increase lifespan, even in the absence of improving known pathology, in the *Cln2*^*R207X*^ mouse model of CLN2 disease.

## Acknowledgements

The authors give a special thanks to Stephen and Sandra Lehrman and the Lehrman Family Fund for providing financial support to complete this work. Thanks also to the Batten Disease Support and Research Association for research support, and to Dr. Russell B. Williams at the Proteomics and Mass Spectrometry Facility of the Donald Danforth Plant Science Center who performed LC-MS/MS analysis on instruments purchased with DDPSC funds.

